# Supression of Perforin-like Protein pores inhibit *Plasmodium* multistage-growth, transmission and erythrocyte senescence

**DOI:** 10.1101/756197

**Authors:** Swati Garg, Abhishek Shivappagowdar, Rahul S. Hada, Rajagopal Ayana, Chandramohan Bathula, Subhabrata Sen, Inderjeet Kalia, Soumya Pati, Agam P Singh, Shailja Singh

## Abstract

The pore forming *Plasmodium* perforin like proteins (PPLP), expressed in all stages of the parasite life cycle are central drivers for host interactions critical for completion of parasite life cycle and high transmission rates. The high sequence similarity in the central membrane attack complex/ perforin (MACPF) domain and consequent functional overlaps defines them as an attractive target for the development of multi-stage antimalarials. Herein we evaluated the mechanism of pan active function of central, highly conserved region of PPLPs, MACPF domain (PMD) and inhibitory potential of specifically designed anti-PMD chemo. The *E. coli* expressed rPMD interacts with erythrocyte membrane and form pores of ~10.5 nm height and ~24.3 nm diameter leading to haemoglobin release and dextran uptake. The treatment with PMD induced erythrocytes senescence at 48 hours which can account for the physiological effect of disseminated PLPs in loss of circulating erythrocytes inducing anemia during malaria infection. The anti-PMD inhibitors effectively blocked intraerythrocytic growth by suppressing invasion and egress of merozoites and protecting against erythrocyte senescence. Moreover, these inhibitors also blocked the hepatic stage and transmission stage parasite development suggesting multi-stage and transmission-blocking potential of these inhibitors. Additionally, the erythrocyte senescence protective potential of PMD inhibitors can be used to occlude PPLPs mediated severe malarial anemia. Further these inhibitors can be developed with a potential to protect against severity of the disease.

**Author Summary:** Malaria continues to be a major global health threat despite of several exciting improvements in the treatment and prevention of the disease. One of the major concerns in the development of therapy is the emergence of the drug resistance. But for the efficient treatment regime, targeting multiple stages including host and vector would serve as an ideal therapy. Perforin like proteins (PLPs) are eukaryotic pore forming proteins that are highly conserved in the apicomplexan parasites. These play crucial roles in entry and exit of parasites from the host cells and establish infection at multiple stages of *Plasmodium spp.* life cycle. Understanding the mechanism of pore formation by smaller, functional, pan-active scaffold of PLPs can serve as a target for development of cross stage protection. Here, using various biochemical, biophysical and pharmacological evidences, we validate the activity and characterize the pore formation of PLPs on erythrocytes. Further, our specifically designed inhibitors could restrict this pore formation and impede the exit/entry of the parasites. Moreover, these inhibitors could also exert multiple stage inhibition and rescue the uninfected erythrocytes from death. Together, this study highlights the mechanism of pore formation by PPLPs and evaluates their potential for the development of pan-active inhibitors to provide both symptomatic and transmission blocking cure for malaria.

## Introduction

Malaria remains a serious global health challenge and major roadblock for the economic growth of the poor and developing economies. The rapid emergence of drug resistant malaria parasites has exceeded the rate at which anti-malarial therapies are presently being introduced. Currently available antimalarial therapies target blood stage with an aim to lower parasite burden and cure the disease. However, the next generation antimalarials, according to the criteria set by WHO, should also provide cross-stage protection to prevent further spread of the disease(1). Therefore, an efficient treatment regimen needs to be both curative and transmission blocking. In lieu of the same, molecular players performing multiple roles across the life cycle of the parasite would thus serve as an ideal target for the development of pan-active therapeutic interventions. *Plasmodium* Perforin like proteins (PPLPs) are excellent candidates in this regard and needs to be further characterised(2).

Perforin like proteins (PLPs) are the eukaryotic pore forming proteins conserved across the apicomplexan parasites(2). They are the crucial players in the biology of malaria parasite across all the stages *Plasmodium* life cycle(3,4). The genome of *Plasmodium spp.* encodes for five PPLPs (PPLP1-5) that work in different combinations at different stages of parasite life cycle and are indispensable for the parasite growth and survival(5,6). In the liver stage PPLP1 has a role in successful establishment of hepatocyte infection(7). PPLP1 and PPLP2 are expressed in blood stage schizonts, merozoites and are involved in permeabilization of host erythrocyte membrane during the egress of malarial parasites(2). They are also proposed to play a role in modulation of host erythrocyte calcium during invasion of merozoites (unpublished data). In the gametocytes, PPLP2 is responsible for egress of activated male gametocytes from the host erythrocytes(8,9). PPLP3, PPLP4 and PPLP5 are expressed in ookinete and are involved in mosquito midgut traversal to form oocysts(10–12). Despite the importance that PPLPs have in parasite life cycle, no chemotherapeutic interventions have been developed against them. The very few reported inhibitors for eukaryotic pore forming identified to date are mostly exerting their effects indirectly through inhibition in protein processing, storage, or secretion from organelles rather than directly inhibiting perforin’s function in the target cell(13,14). Recently, some small molecule inhibitors were identified from a high throughput screen which inhibited mouse perforin at sub micromolar doses (15,16). These studies provide motivational for the development of anti-PPLP molecules but requires, identification and functional characterization of a common motif of PPLPs that can serve as the universal target for chemotherapeutics.

PPLPs have a N-terminal signal sequence, a central MACPF (membrane attack complex/perforin) domain and a C-terminal β-sheet rich domain. The central MACPF domain is the functional unit of PPLPs having the characteristic signature motif of eukaryotic pore forming proteins and two transmembrane helical domains (CH1 and CH2) which exhibit the typical arrangement of alternate hydrophilic and hydrophobic residues and is involved in membrane insertion(17–19). The C-terminal domain (CTD) is rich in β sheet and is involved in membrane binding similar to eukaryotic pore forming proteins(20). PPLP molecules are initially secreted as monomers which bind to the target cell membrane and oligomerize on its surface to form functional, transmembrane pores(18,21). The structure of PPLPs remain unresolved, however, recently the crystal structure of a closely related PLP of another apicomplexan parasite *Toxoplasma gondii*, TgPLP1 was deciphered (22). The structure of TgPLP1 is similar to the reported eukaryotic pore forming proteins. But, it is bulkier than others in terms of the presence of extra helices that may play roles in pore formation (22). The structure of PPLPs can now be predicted using TgPLP1 as template for designing anti-malarial chemotherapeutics.

In this study, we have identified a central, highly conserved region, MACPF domain, of *P. falciparum* PLPs (PfPLPs) and further show that it is a pan-active drug target for multi-stage malaria interventions. The pan-MACPF domains (PMD) of PfPLPs display the conserved pore forming activity which is confirmed by hemoglobin release assay and dextran uptake. To gain insights into mechanical characteristics of pore, AFM of erythrocyte surface having PMD pores was performed which revealed that PMD form pores of 10.5nm height and 24.3nm diameter. We further designed anti-PMD inhibitors (PMIs), and confirmed their pan-inhibitory potential against PPLPs. Binding of PMIs to PMD arrested pore formation on erythrocyte membrane. *Ex vivo*, restriction in pore formation by PMIs impeded parasite egress from and invasion into host erythrocytes. Moreover, PMIs also inhibited gametocyte exflagellation and sporozoite infection to hepatocytes demonstrating the potential to deliver stage-independent protection. Interestingly, we further discovered that at sublethal concentrations, PPLPs induce calcium influx in bystander erythrocytes leading to their premature cell senescence which could be associated with erythrocytic lysis contribution to severe malaria anemia. Taken together, the findings from the present study document the feasibility for the development of pan-active molecules against MACPF domain to provide both symptomatic and transmission blocking for curing malaria and sets the stage for the development of promising chemotherapeutics.

## Results

### The pan-MACPF domain of PfPLPs forms pores at physiological relevance

The presence of MACPF domain in PfPLPs is necessary for pore formation. We mapped the minimal, active domain of PfPLPs and named it as pan-MACPF domain (PMD) (S1A Fig). The PMD harbours a signature motif (Y/W)-X6-(F/Y)GTH(F/Y)-X6-GG) along with two cluster of helices (CH1 and CH2) to form pores in the cell membrane (Fig 1A). Interestingly, the sequence analysis of different PMDs revealed that they are highly conserved and closely related to each other. This stretch of sequence is also evolutionarily conserved across *Plasmodium spp* (S2A Fig).

**Fig 1.**
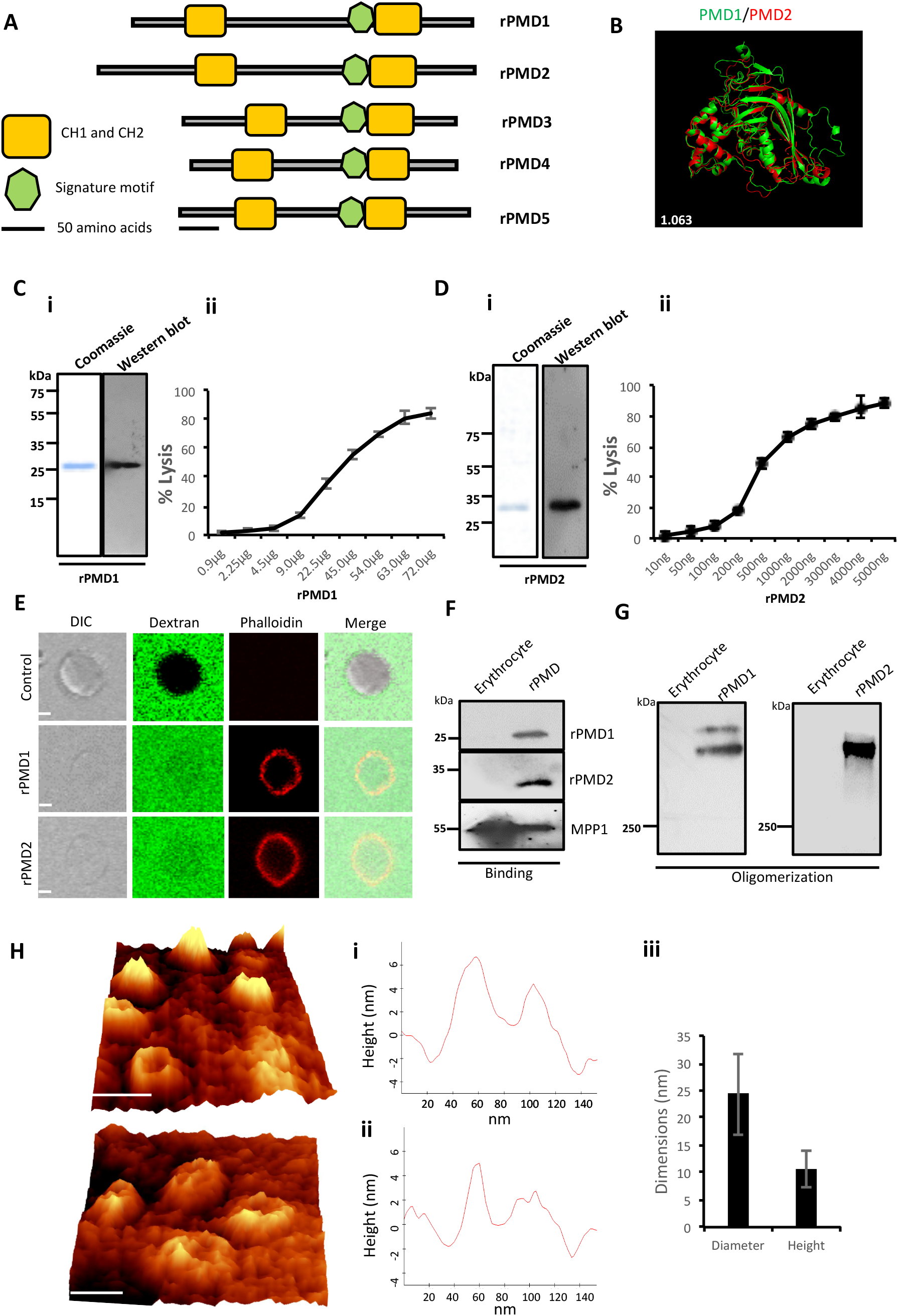
Purification and activity of rPMDs. (**A**) Domain architecture of MACPF domain of PfPLPs. The signature motif (green box) and two transmembrane helical domains, CH1 and CH2 (yellow boxes) are depicted. Scale bar represents 50 aa. (**B**) Structural superimposition of PMD domain of PfPLP1 and PfPLP2. RMSD value is indicated in white. (**C**) (**i**) Coomassie and Western blot of affinity-purified rPMD1 probed with anti-his antibody. (**ii**) Dose-dependent membranolytic activity of rPMD1. Lysis of human erythrocytes was analyzed in the presence of different concentrations of rPMD1. The graph indicates per cent lysis of human erythrocytes by rPMD1 as compared with 100% hypotonic lysis of human erythrocytes in water. (**D**) (**i**) Coomassie and Western blot of affinity-purified rPMD2, probed with anti-his antibody. (**ii**) Dose-dependent membranolytic activity of rPMD2. Lysis of human erythrocytes was analyzed in the presence of different amount of rPMD2. The graph indicates per cent lysis of human erythrocytes by rPMD2 as compared with 100% hypotonic lysis of human erythrocytes in water. (**E**) Permeabilization activity of rPMD1 and rPMD2. The human erythrocytes were incubated with Phalloidin Alexa 594 and 10kDa FITC-Dextran in the presence and absence of rPMD1 and rPMD2 and visualized under confocal microscope. Phalloidin staining was detected in the human erythrocytes treated with rPMD1 and rPMD2 but not in untreated human erythrocytes. (**F**) Western blot analysis of bound rPMD1 and rPMD2 eluted by 1.5M NaCl. (**G**) Oligomerization of rPMD1 and rPMD2. rPMDs are incubated with human erythrocytes at 37°C for 30 mins and analyzed for oligomeric rPMD1 and rPMD2 by Western blotting. (**H**) Visualization of oligomeric pores by AFM. (**i**) Erythrocytes were treated with rPMD2 for 30 mins at 37°C and visualized under the AFM for pore formation. The image was 3D constructed using project Witec 4.1 software. Scale bar represents 50 nm. (**ii**) Line profile of rPMD2 oligomers. Representative images of the height of oligomer measured along the pore in AFM topographs is depicted. (**iii**) Average ring diameter and height of rPMD2 treated oligomers protruding from erythrocytes membrane. Diameter and height were measured for oligomers formed on the erythrocyte membrane. Bars represent average of 30 oligomers and error bars SD.

To investigate whether, the conservation of sequence is also reflected in structural conservation, we *in silico* modelled the structure of PMDs based on MACPF domain of a closely related apicomplexan parasite *T. gondii*, TgPLP1. Like other MACPF domains, PMDs contain a central β-pleated sheet, surrounded by CH1 and CH2 on either side. In addition, two helical inserts are additionally present in PMDs, similar to TgPLP1, that are absent in all other reported structures of MACPF domain. Overall, the structural prediction revealed significant conservation of MACPF domain fold across different PMDs (Fig 1B and S1B Fig).

The expression and purification of recombinant MACPF domain from the earlier reported mammalian or insect cell system yield insufficient quantities of recombinant protein. To characterize, the pore forming activity of PMDs *in vitro*, we cloned MACPF domain of PfPLP1 (rPMD1) and PfPLP2 (rPMD2) in pET28a (+) and recombinantly expressed rPMD1 and rPMD2 in *E. coli*. We could successfully purify the active protein from bacteria (Fig 1C(i) and 1D(i)) and demonstrate the erythrocyte lysis activity of both, rPMD1 and rPMD2, *in vitro* (Fig 1C(ii) and 1D(ii)). Both PMDs lysed erythrocytes in dose dependent manner which is due to their pore forming activity on erythrocyte membrane.

To confirm the permeabilization activity of rPMDs, the activity of rPMD1 and rPMD2 was monitored in the presence of rhodamine Phalloidin and 10 kDa FITC-dextran. The rPMD1 and 2 treated erythrocytes displayed phalloidin positivity accompanied by the uptake of 10 kDa FITC-dextran indicating permeabilization of the cell membrane (Fig 1E).

The PFPs oligomerizes in lipid bilayer following the binding of monomer and create pores. The binding of rPMDs to the erythrocytes was tested by Western blotting and demonstrated that rPMD1 and rPMD2 binds to erythrocytes in monomeric form (Fig 1F). We further investigated whether binding of rPMDs leads to their oligomerization. We found that rPMD1 and rPMD2 formed SDS resistant, higher molecular weight oligomers (>250kDa) suggesting involvement of more than 8 monomers in formation of pores (Fig 1G).

Although, many studies indicate the pore forming capacity of PPLPs, this process has not been characterised in detail. Hence, to gain in-depth insights into the characteristics of PPLP pores, we performed high resolution atomic force microscopy (AFM) of oligomers on erythrocyte membrane. Erythrocytes serve as the physiological host of PfPLP1 and PfPLP2 *in vivo* and hence we performed studies on red cells rather than model lipid bilayer membranes. AFM topographs of erythrocytes incubated with rPMD2 showed pore forming oligomers in circular forms (Fig 1H). Height analysis demonstrated that rPMD2 oligomers have vertical measurement of 10.55±3.22nm (mean ± SD, n=30) on the erythrocyte surface (Fig 1H(i) and 1H(ii)). The diameter of oligomers was distributed widely between 10nm to 40nm, showing a mean at 24.33±7.42nm (mean ± SD, n=30). The line profile reveals the height and diameter of a single pore formed by rPMD2 (Fig 1H(iii)). Overall, this result confirms that PMD can drill pores in the membrane of erythrocytes.

### Anti-PMD inhibitors bind to PfPLPs

The evolutionary conservation of PPLPs and their importance in disease pathogenesis prompted us to develop novel inhibitors against pan-MACPF domain. Recent reports demonstrate the identification of anti-perforin inhibitors through a high-throughput screen(23). Inspired from this, anti-PMD inhibitors (PMI), C01 and C02, were designed with a hypothesis that they can be pan-active against all PPLPs (Fig 2A). To validate the binding of PMIs to PMDs in silico docking analysis was performed. The structure-refined models of PMD1-5 having a RMSD score of <2.0 were used for docking with PMIs. The molecular docking results revealed that C01 and C02 could efficiently bind to PMD1-5 with similar docking energies (Fig 2B (i-ii), Fig 3C and S3B (i-iii) Fig). The similar binding energy suggests that PMIs can inhibit PMDs with identical activity *in vitro*. *In silico* data further revealed that PMIs are binding in the same pocket of all PMDs. This pocket is not only structurally conserved but also has sequence identity as revealed by sequence alignment of PfPLP1-5 (S3A Fig).

**Fig 2.**
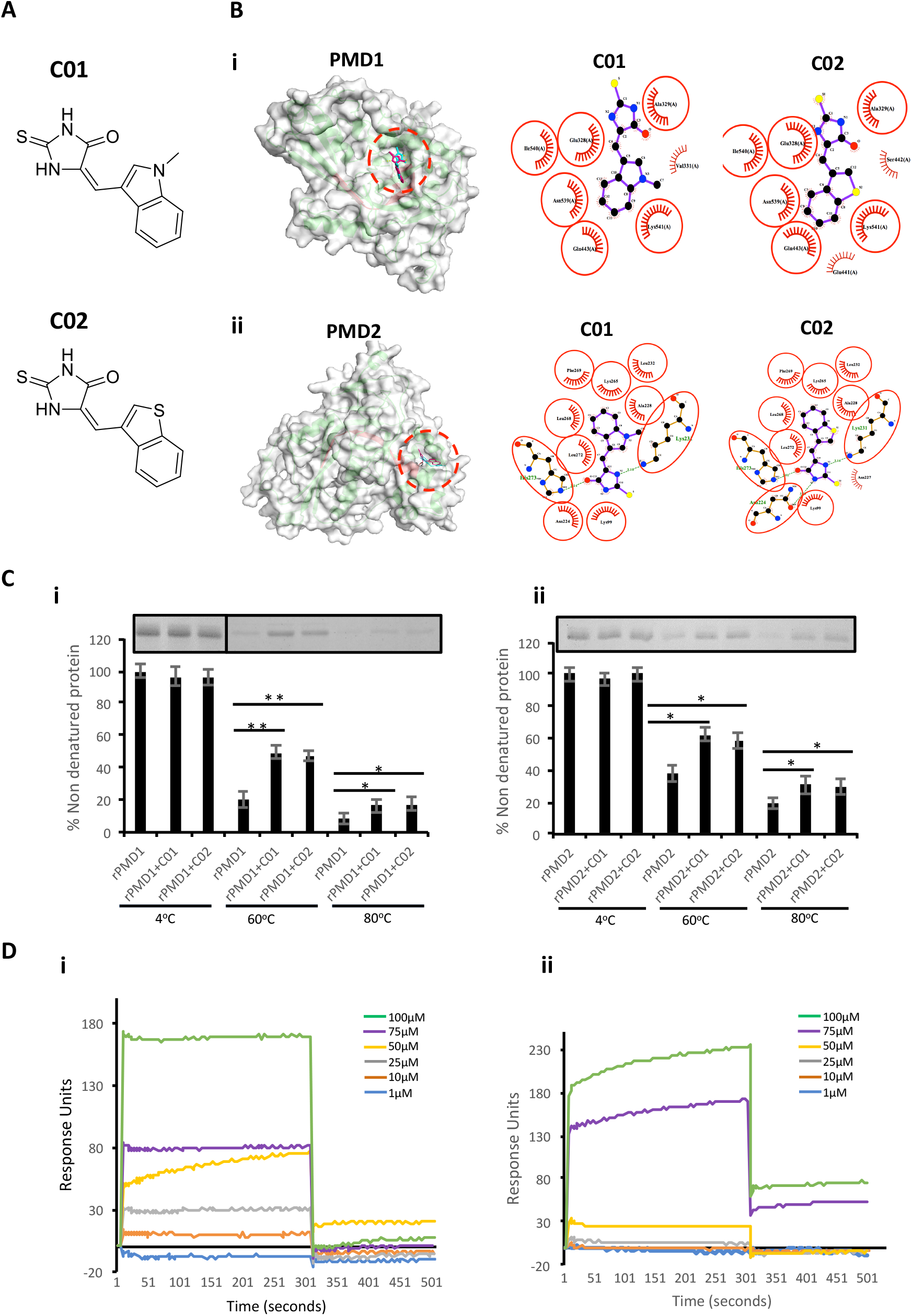
*In silico* and *in vitro* interaction of C01 and C02 with rPMDs. (**A**) (**i**) The structures depict scaffold of (Z)-5-((1-methyl-1H-indol-3-yl)methylene)-2-thioxoimidazolidin-4-one (C01) and (**ii**) (Z)-5-(benzo[b]thiophen-3-ylmethylene)-2-thioxoimidazolidin-4-one (C02). (**B**) The surface images of docked complexes have shown efficient binding of C01 and C02 to the signature motif of PMD1 (**i**) and PMD2 (**ii**). The ligplot figures demonstrate the specifics of the atoms involved in interaction. (**C**) Interaction of PMIs and PMDs by CETSA. The drug target engagement between the compounds and recombinant proteins was analysed by subjecting the samples to thermal denaturation at 60°C and 80°C (**i**) (**ii**). The protein intensity at 4°C was taken as control. The band intensities graph was plotted considering the 4°C samples as 100% non-denatured protein. Error bar represents SD (p<0.05*; p<0.01**). (**D**) rPMD1 was immobilized onto a nickel charged NTA SPR chip. C01 (i) and C02 (ii) were injected over immobilized rPMD1. The PMIs show concentration dependent binding to rPMD1.

**Fig 3.**
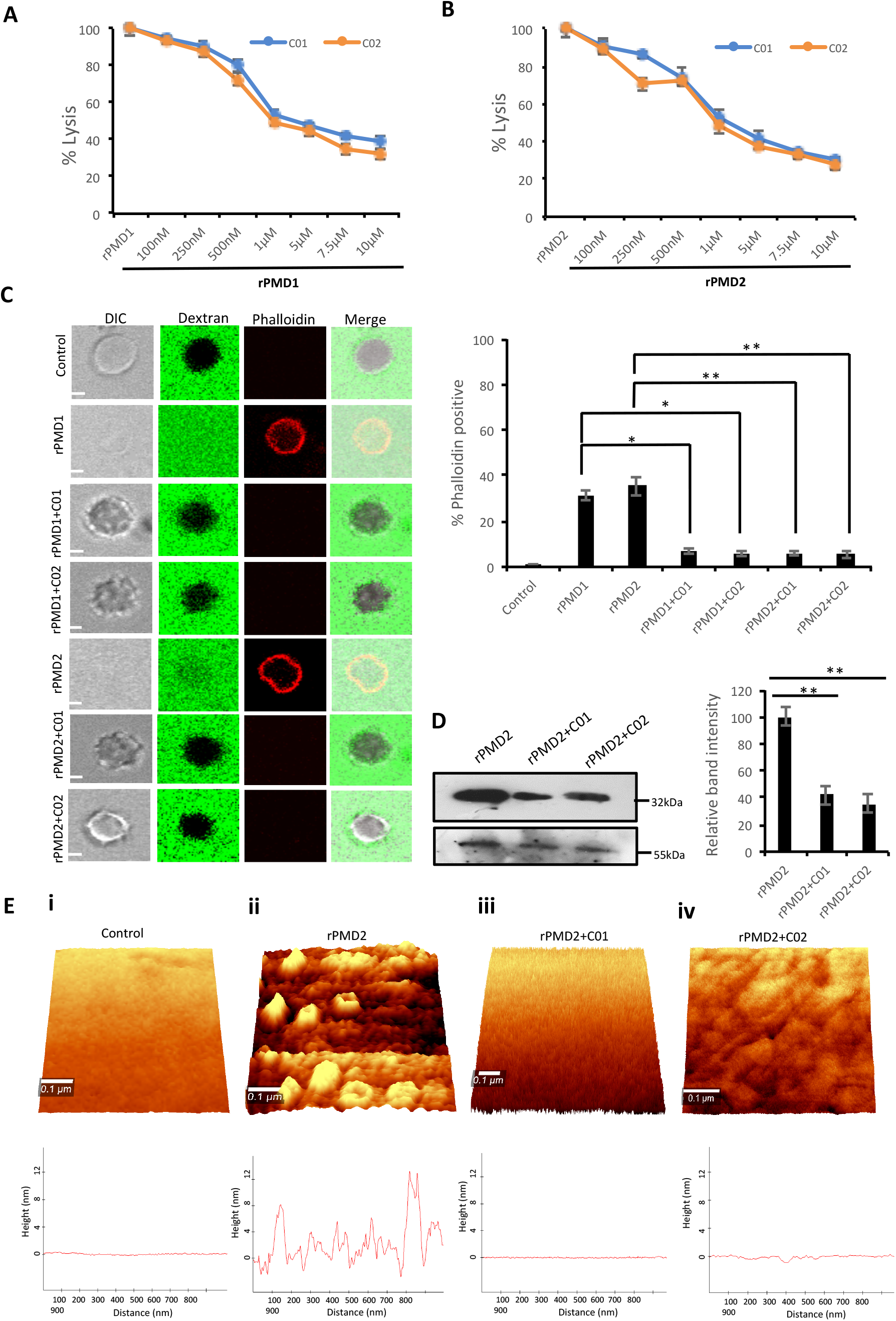
C01 and C02 rescue RBCs from rPMD mediated lysis. (**A**) Anti-PLP effect of C01 and C02. The inhibition in erythrocytes lysis was evaluated by treating different concentration of compounds with IC_50_ of rPMD1 (**A**) and rPMD2 (**B**). The lysis caused by recombinant protein alone was considered as 100% and was plotted respectively for the compound treated erythrocytes. Error bar represents SD. (**C**) Inhibition of permeabilization activity by PMIs. The erythrocytes were incubated with Rhodamine-phalloidin and 10kDa FITC-dextran in the presence of recombinant proteins in presence or absence of PMIs and visualized under the confocal microscope. PMI treatment inhibited phalloidin staining and dextran entry into the erythrocytes. The phalloidin positive erythrocytes were counted under the microscope for different treatments and their relative percentage was plotted. Error bar represents SD (p<0.05*; p<0.01**). (**D**) Inhibition of PMD binding to erythrocyte membrane in presence of PMI. Western blot was performed to detect the effect of PMIs on the binding of rPMD2 to the erythrocyte membrane. MPP1 was taken as the initial erythrocytes loading control. The relative band intensity of bound rPMD2 was analysed using ImageJ. Error bar represents SD (p<0.01**). (**E**) Erythrocytes were treated with rPMD2 in the presence and absence of PMIs and pore formation was observed under the AFM. (**ii**) The PMD treated erythrocytes demonstrate formation of pores on erythrocyte surface. The line profile further depicts the roughness of erythrocytes. The PMD treated erythrocytes (**iii**, **iv**) in the presence of PMIs does not demonstrate formation of pores and the surface roughness was also reduced to normal erythrocytes (**i**).

To evaluate the binding of PMI to the rPMD, we modified and performed cellular thermal shift assay (CETSA). This technique involves the detection of target protein by monitoring the thermostability of native protein in presence of its selective inhibitor. In principle, specific binding of the drug to its target protein increases the stability of protein at high temperatures(24). In similar line, we treated the purified rPMD1 and rPMD2 with C01 and C02 and heated them at 60°C and 80°C while the sample at 4°C, served as loading control. Analysis of band intensities demonstrated significant thermal protection of both rPMD1 and rPMD2 in the presence of compounds suggesting that C01 and C02 are interacting with rPMDs (Fig 2C (i-ii)). Further, the interaction of PMIs to PMDs is strong enough to impart thermal protection to the recombinant protein.

Surface Plasmon Resonance (SPR) is a sensitive technique to validate the protein-protein as well as protein-drug interactions. To further confirm the interaction of PMIs with rMAC1, we performed SPR with different compound concentrations (1-100 μM). In both the cases, the compounds showed concentration dependent binding to the coated rMAC1 (Fig 2D (i-ii)). Together, SPR and CETSA confirms the binds of PMIs to rPMD.

### PMIs inhibit PMD mediated pore formation

We analysed the inhibitory activity of PMIs by performing rPMD mediated erythrocyte lysis assay in the presence of C01 and C02. The results demonstrated that C01 and C02 could inhibit activity of both rPMD1 and rPMD2 in sub-micromolar range (Fig 3A and 3B). Together, these data demonstrate the designing of anti-PMD inhibitors that inhibits the pore forming ability of PMDs.

To investigate complete disruption of pore formation in PMI treated erythrocytes, influx of small molecules such as rhodamine-Phalloidin (~1kDa) and FITC-dextran (10kDa) was investigated. PMD treated erythrocytes could not uptake dextran or Phalloidin in presence of PMI as compared to PMI treated erythrocytes (Fig 3C). This confirms that there is complete abrogation of pore formation in presence of PMIs.

Since the pore formation is chiefly dependent on binding of monomers, we tested binding of monomer to erythrocyte membrane in the presence of PMIs. rPMDs could not bind to erythrocyte membrane in the presence of PMIs suggesting that inhibition in pore formation is due to restriction of monomer binding to erythrocyte membrane (Fig 3D). Next, to closely monitor that PMIs are completely abrogating formation of pores and there is little ot no change in membrane roughness due to rPMD binding, AFM was performed to evaluate surface topology of treated erythrocytes. We could clearly detect formation of pores on the erythrocyte membrane treated with rPMD2 (Fig 3E (ii)). Also, the surface of rPMD treated erythrocytes was very rough and uneven. In comparison, treatment with rPMD2 in presence of PMIs, displayed no evident structures on the surface of erythrocytes, indicating the impairment in oligomerisation and subsequent pore formation (Fig 3E (iii-iv)). Further the smooth surface of PMI treated erythrocytes, similar to untreated erythrocytes, suggest that PMIs could revert all the phenotypes induced by rPMDs (Fig. 3E (i-iv)). Taken together, this data indicates that C01 and C02 inhibit pore formation by a decrease in binding of the rPMD to the erythrocyte membrane.

### PMIs demonstrate anti-parasitic activity at multiple stage of parasite life cycle

To monitor the effect of PMIs on parasite growth, Giemsa stained smears were prepared for the treated as well as untreated parasites at different time points. Drug treatment did not affect the parasite growth from ring to schizont stages. However, the treated parasites could not egress, and the merozoites that came out, could be seen attached to erythrocytes but not forming rings (Fig 4A). Having confirmed the inhibitory activity of PMIs we further evaluated their anti-malarial activity. The ring infected parasites (~5-6hpi) were treated with different concentrations of drug for 72h. The parasites treated with DMSO served as a control and the parasite growth was measured by fluorimetry using SYBR green I-based assay. Both C01 and C02 demonstrated the ability to inhibit parasite growth with an IC_50_ of 3.54 μM for C01 and 3.31 μM for C02 (Fig 4B). This is similar to the reported role of PfPLPs in egress and invasion of merozoites suggesting the specific action of PMIs towards PMDs.

**Fig 4.**
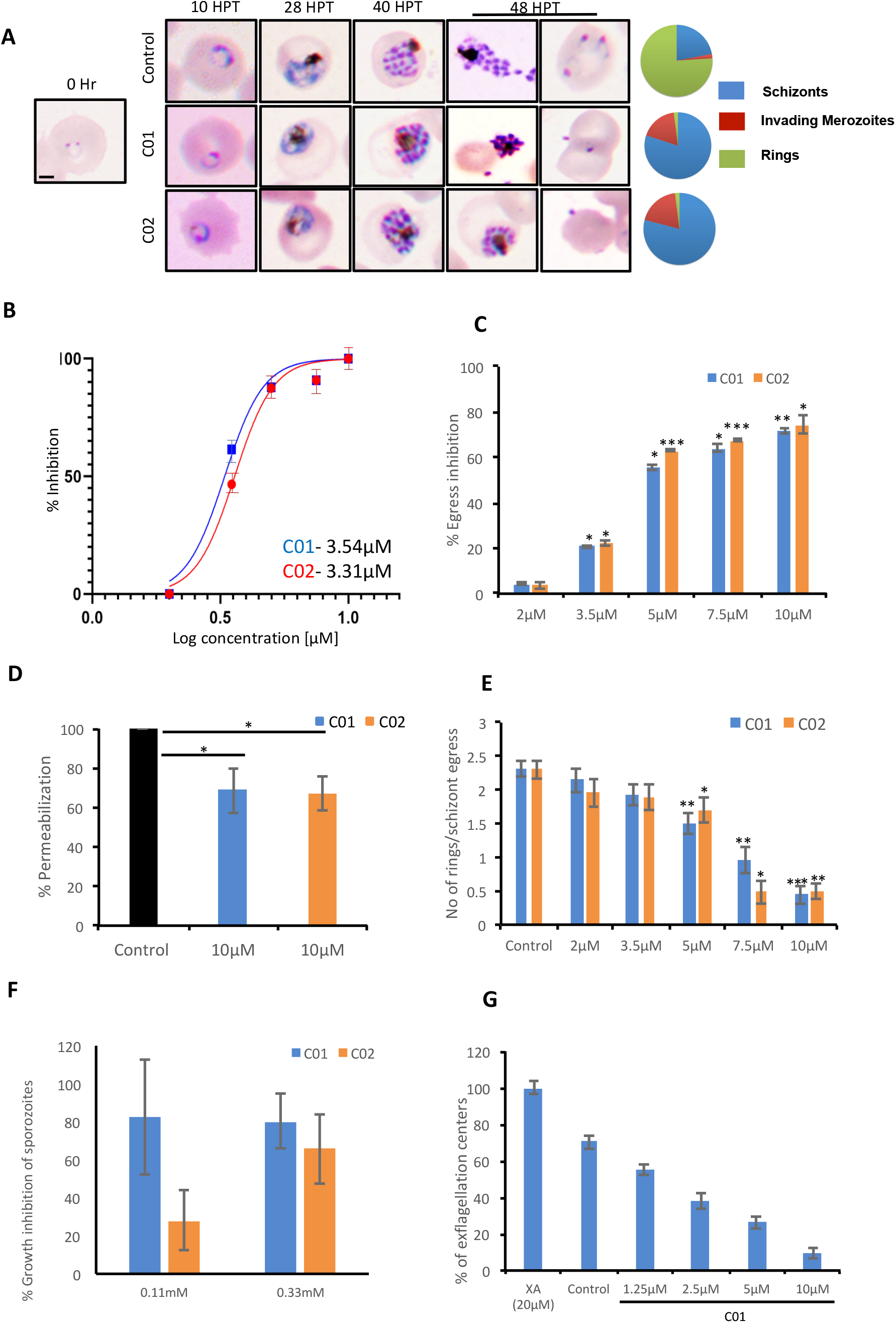
C01 and C02 show multistage inhibition. (**A**) Images of Giemsa stained smears of PMI treated and untreated parasites. Early rings were treated with PMIs and smears were prepared 10, 28, 40 and 48 h post treatment (HPT). Pie charts show relative proportions of schizonts, invading merozoites and rings. (**B**) C01 and C02 inhibit the *in vitro* growth of *P. falciparum*. The effect of PMIs on growth inhibition of *P. falciparum* was evaluated after one cycle of parasite growth. Scale bar represents 2 μm. (**C**) Late stage schizonts were treated with different concentrations of compounds. The relative inhibition in egress was calculated by counting the number of remaining schizonts after 7 hrs of treatment as compared to control. Error bar represents SD (p<0.05*; p<0.01**; p<0.005***). (**D**) Permeabilization of PMI-treated schizonts was measured by flow cytometry using Phalloidin. Error bar represents SD (p<0.05*). (**E**) Late stage schizonts were treated with different concentrations of PMIs. The ability to form rings per schizont egress was calculated by counting the number of schizonts and rings after 7 hrs using giemsa stained smears. Error bar represents SD (p<0.05*; p<0.01**; p<0.005***). (**F**) Treatment of male gametocytes with PMIs inhibits exflagellation. The number of exflagellation centres was scored 15 minutes post activation in 30 optical fields at 40X magnification by light microscopy. Exflagellation efficiency of PMI treated versus untreated gametocytes is shown. (**G**) HepG2 cells were infected with *P. berghei* sporozoites and treated with PMIs. Parasite growth was assessed after 51 hours via real-time PCR using *P. berghei* 18S rRNA specific primers. Parasite growth inhibition was calculated by dividing the 18S rRNA copy number of the experimental group by that of the untreated control group. The fraction obtained was then converted into % inhibition (with respect to untreated as 100%).

To confirm the egress defect, late schizonts (~44-46hpi) were treated with different concentrations of C01 and C02 for 6 hrs and decrease in number of schizonts was scored (Fig 4C). The intra-cellular Ca^2+^ chelator BAPTA-AM was used as a positive control. We observed a dose dependent defect in egress. PfPLPs have a role in host erythrocyte permeabilization that facilitates egress of merozoites. We observed that PMI treated schizonts demonstrated restriction in permeabilization of host erythrocyte as compared to untreated schizonts (Fig 4C and 4D), implying specific action of PMIs towards PfPLPs in egress.

Since, the giemsa scoring also suggested a defect in invasion, we scored for ring formation after 10 h post-treatment. To specifically assess the effects of C01 and C02 on invasion and avoid the effects of their influence on egress, we calculated number of rings formed per schizont egress. A drastic decrease in the number of rings formed in treated parasites as compared to the untreated control was noted (Fig 4E).

To rule out the possibility that drug treatment has any effect on ring and trophozoite stages, we devised a novel ring toxicity assay and trophozoite toxicity assay. In these assays, rings or trophozoites were treated for 6 h, washed and accessed for their growth during the next cycle. If the compounds have any toxicity on these stages, it will be reflected in this assay. Our data suggests that these compounds do not exhibit any toxicity towards either ring and trophozoite stages (S4A and S4B Fig). Taken together these findings imply that both compounds act only on the stages which involves the PPLP activity and can be used as generic inhibitor of PPLPs. Since PPLPs play a role during multiple stages of parasite cycle, we studied their cross-stage inhibitory potential. PPLP1 plays a role in sporozoite infection to HepG2 cells. To test the inhibitory effect of PMI on sporozoite infection, we infected HepG2 cells with *P. berghei* sporozoites treated with different concentration of PMI for 72 h. The sporozoite infection was scored using real-time PCR analysis. As shown (Fig 4F), both compounds displayed an inhibitory effect toward *P. berghei* sporozoites. Both the compounds showed no defect in viability of HepG2 cells suggesting the specific effect towards sporozoite but not host (S4C Fig).

PPLP2 plays a role in gametocyte egress and hence the effect of PMI was monitored during this stage. The egress efficiency of male gametocytes, as revealed by counting exflagellation centres, was markedly reduced following PMI treatment (Fig 4G). Together, this suggests the cross-stage anti-malarial activity of PMIs which can be further exploited to deliver both symptomatic and transmission blocking cure for the disease.

### Sub lytic concentration of PMDs trigger premature senescence of bystander erythrocytes

PfPLP1 and PfPLP2 are secreted during egress of merozoites and binds to the erythrocyte membrane(2). To analyse the role of secreted PfPLPs, time lapse microscopy was performed, mimicking *in vivo* condition. Addition of sublytic concentration of rPMD1 and rPMD2 induced calcium influx in erythrocytes leading to changes in erythrocyte deformability followed by echinocytosis (Fig 5A, S1 Movie). Inhibition of PMD binding to erythrocyte by PMIs restricted the influx of calcium and echinocytosis (Fig 5A, S2 Movie, S3 Movie). The calcium intake was further quantitated by flow cytometry that also showed increase in intracellular calcium in response to rPMDs. This result suggests that PfPLPs form smaller pores on erythrocytes at sub lytic concentrations that induces calcium influx but does not leads to lysis of erythrocytes.

**Fig 5.**
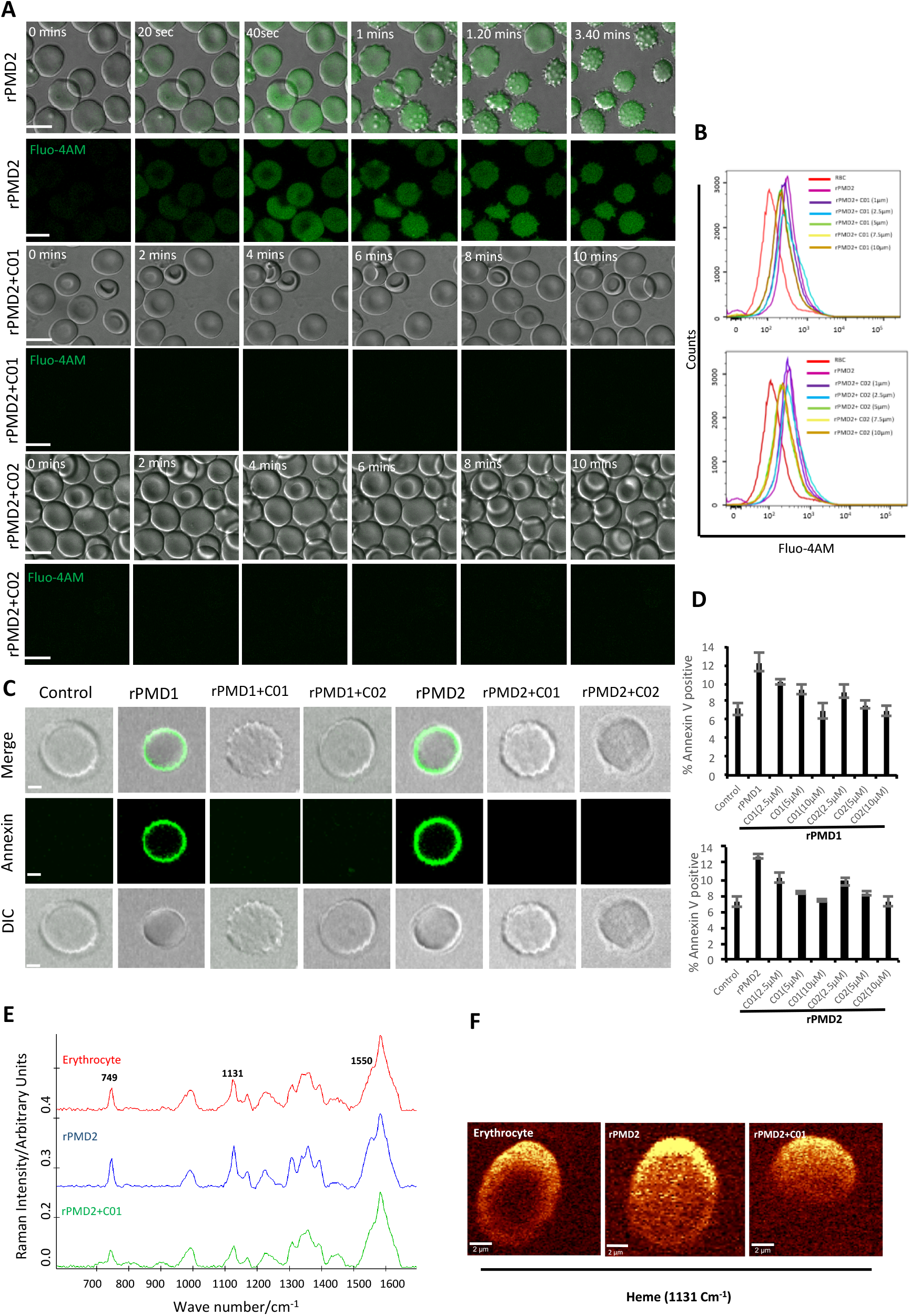
Inhibition of by stander effect mediated by C01 and C02. (**A**) Time course of rPMD2-induced calcium channel formation. Erythrocytes were loaded with Fluo4-AM. Sub lytic concentration of rPMD2 was added and increase in calcium was monitored by confocal microscopy. Selected pairs of DIC and fluorescence images with time elapsed between frames in seconds (Sec) are shown. In case of PMI treatment, C01 or C02 were added along with rPMD2. (**B**) PMI mediated inhibition in calcium influx was measured by flow cytometry. The Fluo-4AM loaded erythrocytes were treated with rPMD2 in the presence and absence of indicated concentration of PMIs and calcium increase was monitored by Flow cytometer. (**C**) Phosphatidylserine exposure on erythrocyte surface. The erythrocytes were treated with rPMD1 or rPMD2 in the presence or absence of PMIs and stained with Annexin V-FITC after 48 hrs. The stained erythrocytes were visualized under confocal microscope. (**D**) The annexin positive erythrocytes were quantitated using flow cytometer. (**E**) Average Raman spectra of the untreated erythrocytes or erythrocytes treated with rPMD2 in the presence of absence of C01 was captured using 532 nm excitation. All the raman spectra were presented after pre-processing (baseline correction, smoothening and background removal) using asymmetric least squares smoothing method. (**F**) Raman images of erythrocytes was observed at 1131cm^−1^ which demonstrates the distribution of methemoglobin.

Calcium influx also leads to phosphatidyl serine (PS) exposure on surface of erythrocytes that can lead to phagocytosis of cells by macrophages. To assess this effect, erythrocytes were treated with sub lytic concentrations of rPMD1 and rPMD2 in the presence and absence of PMIs and stained with Annexin V-FITC after 48 h. The Annexin V-FITC positivity was analysed by microscopy and flow cytometry (Fig. 5C,S4D Fig). rPMDs treated erythrocytes in absence of PMIs were Annexin-positive suggesting induction of PS exposure while the presence of PMIs abrogated their Annexin positivity (Fig 5C (i-ii)). This result indicates that at sub lytic concentrations, PfPLPs can induce delayed erythrocyte sequestration.

Oxidation of hemoglobin, leading to methaemoglobin formation, is a marker of erythrocyte senescence. The methaemoglobin caries the oxidized form of heme (Fe^3+^) that leads to changes in its porphyrin ring(25). To detect these changes in heme, Raman spectroscopy was used. Raman spectroscopy is a non-invasive and label free technique for detection of metabolic changes at single cell level. When 532nm is applied to rPMD2 treated erythrocytes, *v*_15_ gains intensity (pyrrole gains intensity) which is co-related well with increase in intensity at band 749cm^−1^. A similar increase of band intensity was observed for band located at 1131cm^−1^ that suggests the asymmetrical pyrrole half-ring stretching vibration (Fig 5D). The 1550 cm^−1^ demonstrates the stretching mode *v* (CbCb)(26). The untreated and PMI treated erythrocytes demonstrated similar peak profile (Fig 5E). Further, Raman imaging of 1131cm^−1^ also confirmed increase in methemoglobin concentration within erythrocytes. (Fig 5F). These results suggest formation of methemoglobin following rPMD2 treatment that could lead to erythrocyte senescence.

## Discussion

Despite abundant evidence that PPLPs are crucial for life cycle progression of malaria parasites, development of chemotherapeutic interventions against them have not been explored. In the blood stage of *P. falciparum*, PfPLP1 and PfPLP2 create pores on erythrocyte membrane that help in the entry (unpublished data) and exit of merozoites(27). In this study, we further propose that the synchronous secretion PfPLPs in blood plasma due to co-ordinated rupture of schizonts *in vivo* leads to formation of pores on bystander erythrocytes leading to their premature senescence and identifies it as one of the probable causes for anemia in severe malaria. This phenomenon was similar to earlier reports where it has been shown that binding of some parasite proteins to erythrocytes leads to anemia(28,29). But the mechanistic role behind the induction of anemia remain to be fully deciphered. However, our data defines that the smaller pores created by PfPLPs induces calcium influx (Fig 5A) in erythrocytes leading to phosphatidylserine exposure (Fig 5C), as signal for recognition by macrophages. This is in line with earlier studies that have demonstrated that increase in intracellular calcium induces premature senescence of erythrocytes(30,31).

The enhanced oxidative stress in erythrocyte senescence is represented by the formation of oxidised form of hemoglobin, i.e. methemoglobin (Fig 5E and 5F)(25,32). The increased levels of methemoglobin *in vivo*, is toxic to the plasma lipoproteins, endothelial cells and other vascular organs. Therefore, the erythrocytes with increased methemoglobin concentrations are cleared from the body through spleen which is a cause of anemia in methemoglobinomia(25,33). The presence of Fe^3+^ heme form in methemoglobin causes changes in pyrrole ring breathing and asymmetrical pyrrole half ring stretching vibrations(26). Using Raman spectroscopy, we could detect the presence of these metabolic signatures in erythrocytes after 48 hours of PfPLP treatment representing the formation of methemoglobin (Fig 5E). The earlier clinical reports advocate the identification of elevated levels of methemoglobin in severe malaria and directly correlates it with the death of patients(34). Our study supports the role of PfPLP in inducing premature senescence of erythrocytes leading to anemia in severe malaria.

The PfPLPs helps in the establishment of parasite infection through different stages of its life cycle. For the development of pan-active anti-PLP therapeutics, we first identified central, pan-active motif of PfPLPs, PMD domain, that is highly conserved across all *Plasmodium* spp (S2A Fig) and oligomerize on host membranes to create pores(18,21). To visualize pore formation at physiological relevance, AFM was performed with modified protocol to overcome the limitations associated with non-adherent, rough cell surface of erythrocyte. The modified protocol could successfully demonstrate that rPMD2 drilled pores of 10.55nm±3.22 nm height and 24.33±7.42 nm width on the erythrocytes (Fig 1H). The characteristics of pore formed by PfPLP are similar to earlier reported eukaryotic PFPs which form smaller pores as compared to prokaryotic PFPs which form larger pores(35). The three-dimensional topology not only confirmed the presence of well-defined pores, but also revealed that the rPMD2 treated erythrocyte surface is much rougher and more uneven as compared to the untreated erythrocytes (Fig 1H (i-ii)). The roughness of erythrocytes reflects its innate response towards any non-physiological, toxic agent which in this case is rPMD2. This information about cellular response is usually missed in other studies where they studied pore formation on model membranes.

Given the multi-step process of pore formation by PLP, there exist multiple stages at which a PLP can be targeted for inhibition. The designed PMIs strongly bind to the PMD domain and inhibits its pore forming activity by restricting the initial attachment of the monomer to the erythrocyte surface (Fig 3D). The decreased binding of monomer inhibits the oligomerization of PMD to form pores confirming that PMIs completely abrogated the activity of PfPLPs. Ring stage parasites treated with PMIs progresses to schizonts without any growth defect, but the mature parasites could not egress and re-invade to new erythrocyte (Fig 4A, 4C and 4D). This is in line with the role of PfPLPs in schizont egress and merozoite invasion and suggests that PMIs are not showing any off-target effect *in vivo*. Furthermore, PMIs inhibited gametocyte egress and sporozoite infection to HepG2 cells validating their potential in blocking transmission of malaria parasites (Fig 4F and 4G). The invasion inhibitory activity of PMIs against sporozoite of *P. berghei* is suggestive of their cross-species activity due to evolutionary conservation and high similarity of PMDs. These scaffolds could be improved for their efficacy and development of better anti-malarials.

Taken together, we identify novel pharmacological inhibitors against PPLPs that can preserve the cell membrane integrity and viability of all host cells and thus aid in facilitating parasite elimination by preventing further infection. Also, the proposed therapy can be used as anti-virulence therapy along with routine medication for improving treatment outcomes. Since the anti-virulence strategies do not focus on directly killing the pathogen, parasite is not under the selective pressure of resistance development. Therefore, our study lends weight for development of novel pharmacological approaches against malaria to not only cure the disease but also ameliorate disease pathogenesis in clinical conditions.

## Methods

### Expression and activity of rPMD1 and rPMD2

Codon optimized gene encoding for rPMD1 and rPMD2 domain, were subcloned into bacterial expression vector PET28a(+) and protein expression in *E.coli* cells was induced with 1mM isopropyl-β-D-thiogalactoside (IPTG). His-tagged rPMD1 was purified from inclusion bodies while his-tagged rPMD2 was purified from the soluble fraction using Ni-NTA chromatography. The concentration of purified protein was measured using BCA estimation kit (Pierce, USA).

For the lysis assay, 5*10^6^ human erythrocytes were incubated with rPMD1 or rPMD2 in lysis buffer for 1h at 37°C as described previously (2). The release of haemoglobin into the supernatant was estimated by measuring absorbance at 405 nm. For the permeabilization assays, lysis was performed in the presence of Rhodamine-phalloidin and 10kDa FITC-dextran.

Binding of rPMD1and rPMD2 to human erythrocytes was performed as described previously (36). Briefly, erythrocytes were incubated with rPMD1 or rPMD2 for 1h at 4°C and bound proteins were eluted using 1.5M NaCl and detected by Western blotting. For the oligomerization assay rPMD1 or rPMD2 was incubated with human erythrocytes for 1 h at 37°C and centrifuged. The oligomerised protein was detected in erythrocyte pellets using Western Blot analysis (2).

For the study of PMD mediated increase of intracellular calcium, erythrocytes were stained with Fluo-4 AM (Life Technologies, USA) treated with sublethal concentration of rPMD1 or rPMD2 for 1 h at 37°C. The samples were imaged under the Nikon A1R microscope to check for the increase in Ca^2+^ levels. The staining was quantified using BD Fortessa (Becton & Dickinson, USA) by scoring 100,000 cells per sample and analysed using FlowJo software (Tree Star Inc, Ashland).

For the inhibitor screening, C01 and C02 were added during the assay at the desired concentration.

### Modified Cellular Thermal Shift Assay (CETSA)

Interaction between the PMIs with rPMDs was tested using CETSA as described previously (24). Briefly, the rPMDs alone or in combination with the compounds were treated at 4°C, 60°C and 80°C and cooled down followed by centrifugation. The supernatant was analysed by SDS-PAGE. The band intensities for each of the protein lane was determined using Image J software (NIH, USA) and plotted considering the band intensity of untreated 4°C erythrocytes as maximum (100%).

### *In vitro* culture of *P. falciparum*

Laboratory strain of *P. falciparum*, 3D7 was cultured in RPMI 1640 (Invitrogen, USA) supplemented with 27.2 mg/L hypoxanthine (Sigma Aldrich, USA) and 0.5% Albumax I (Invitrogen, USA) using O+ erythrocytes in mixed gas environment (5% O2, 5% CO2 and 90% N2) as described previously (Trager& Jensen, 1976). Parasites were synchronized by sorbitol selection of rings and percoll selection of schizonts.

### Cytotoxicity assay

Human liver hepatocellular carcinoma cell line (HepG2 cells) were cultured in Dulbecco’s Modified Eagle’s Medium as described previously(37). HepG2 cells were seeded at a density of 30,000 cells per well and allowed to grow overnight. Adhered cells were treated PMIs and kept for 48 hours. Cytotoxic effect was assessed using in MTT (3-[4,5-dimethylthiazol-2-yl]-2,5-diphenyl tetrazolium bromide) assay (Sigma- Aldrich, USA).

### Parasite Growth Inhibition Assay (GIA)

To access the effect of PMIs on parasite growth, synchronized trophozoites were treated with different concentrations of C01 or C02 compounds along with DMSO control at 37°C for one cycle of parasite growth. The smears were prepared after 48 h and stained with Giemsa solution (Sigma, USA) and scored for infection under light microscope. IC_50_ was calculated using graph pad prism 8.0 (CA, USA).

### Parasite Egress and Invasion Assay

To assess the effect of PMIs on parasite egress, late stage schizonts (~46 hrs post invasion (hpi)) were diluted to a final hematocrit of 2% and parasitemia of ~10% and treated with different concentrations of C01 and C02 along with DMSO control for 6 hours. Subsequently, the erythrocytes were smeared and counted for schizonts as well as rings by staining with giemsa under the Olympus (CH2Oi) light microscope. % Egress was calculated as the fraction of schizonts ruptured in treatment and control during the incubation time as compared with the initial number of schizonts at 0 h, using the formula as described below. % Egress was then plotted considering fraction of schizonts ruptured in control as 100% egress.

% Egress= 100[(I-T)/(I-C)]

I, initial no. of schizonts; T, no. of schizonts in treatment; C, no. of schizont in DMSO control.

For scoring invasion, number of rings formed per egress of schizont was counted and plotted.

### Ring and trophozoites toxicity assay

To assess the effect of C01 and C02 on ring stage, early rings (~6 hpi) at a parasitemia of 1% were treated with different concentrations of PMIs along with DMSO control. All the wells were washed after 6 h and replenished with media and harvested after 48 h. The parasite growth was assessed by fluorimeter using SYBR green (Thermo Fisher Scientific, USA) staining. For trophozoites toxicity assay, early trophozoites instead of rings were taken and similar protocol was performed.

### *In vitro* growth inhibition assay for liver-stage parasites

The diluted PMI solutions were added to 24-well culture plates (final DMSO concentration was 1%) containing human HepG2 cells seeded a day prior to the experiment and 0.5 ml complete Dulbecco modified Eagle medium (DMEM) containing 10% fetal bovine serum (FBS), together with an antibiotic antimycotic. Infection was initiated by adding 10,000 *Plasmodium berghei* ANKA sporozoites. Infected cultures were then allowed to grow at 37°C in a 5% CO_2_ atmosphere for 51 h. Culture medium was changed 24 h after infection, and fresh compounds were added at the same concentration as on previous day, to maintain inhibitor pressure throughout the growth period. At the end of the 51-h incubation period, total RNA was extracted using TrizolTM (Invitrogen, USA) reagent. Reverse transcription from 1 μg RNA was performed using a cDNA synthesis kit (GCC Biotech, India) to obtain cDNA. In a real-time PCR mix (H-eff qPCR mix, GCC Biotech, India) of 20 μl, a cDNA equivalent of 0.1 μg RNA was used. The real-time PCR mix also contained *P. berghei* 18S rRNA specific primers. Real-time PCR was performed using an Eppendorf Mastercycler realplex4, and the copy numbers were calculated, using the known amount of plasmid standard having the amplification target sequence. Parasite growth inhibition was calculated by dividing the 18S rRNA copy number of the experimental group by that of the untreated control group. The fraction obtained was then converted into % inhibition (with respect to untreated as 100%).

### Annexin and calcium staining assays

Erythrocytes were treated with sublethal concentration of rPMD1 and rPMD2 along with different concentration of C01 and C02. The samples were incubated for 48 h at 37°C and 5% CO_2_. After 48 h, the samples were stained with Annexin-FITC (Life Technologies, USA) and Fluo-4AM (Life Technologies, USA). The erythrocytes were imaged using a Nikon A1R microscope and also quantified using FACS BD Fortessa (Becton & Dickinson, USA) using Cell Quest software by scoring 100,000 cells per sample. Samples were analysed using FlowJo software (Tree Star Inc, Ashland) by determining the proportion of FL-2-positive cells in comparison to the stained untreated erythrocytes.

### Gametocyte exflagellation assay

To initiate *P. falciparum* gametocyte cultures, synchronized asexual blood stage cultures were grown to a parasitemia of 10-15%, treated with 50 mM *N*-acetyl-D-glucosamine containing medium for four days to remove asexual stages and maintained in complete RPMI/HEPES to allow gametocyte development. Stage V gametocytes were incubated with activation medium (100 nM xanthuneric acid (XA), 20% AB+ human serum in RPMI1640/HEPES) in the presence of C01 and C02 at 25°C. Activated gametocytes were spread on glass slides 5 minutes after activation, fixed with methanol and stained with Giemsa (RAL Diagnostics, France). Around 200-250 gametocytes were scored to determine the percentage of rounded up gametocytes. The number of exflagellation centers were scored in gametocyte cultures by light microscopy at 40X magnification in 30 optical fields 15 minutes after activation.

### Surface Plasmon Resonance

To determine the C01 and C02 interaction with rMAC1, SPR was performed using Auto Lab Esprit SPR. rMAC1 (10 μM) was immobilized on the surface of nickel charged NTA SPR chip. Interaction analysis was studied by injecting C01, C02 along with rPMD1 over the chip surface, with association and dissociation time of 300 and 150 s, respectively. HEPES buffer was used both as immobilization and binding solutions. Surface of the sensor chip was then regenerated with 50 mM NaOH solution. Data were fit by using Auto Lab SPR Kinetic Evaluation software provided with the instrument.

### *In Silico* docking

Using I-Tasser and PHYRE2, domain-based modelling and threading was performed to obtain the 3D structures of *P. falciparum* PLP1-5. Following this, structure refinement was done using ModRefiner. To characterize the catalytic pocket, SiteHound software was used while utilizing the structural motif of MACPF domain in each of the proteins (PLP1-5). Further, molecular docking of two PMIs (C01 and C02) was done using AutodockTools and Autodock vina with exhaustiveness cut-off of 8 for each run. All the ligands were kept flexible by examining torsions.

### Atomic force Microscopy

Erythrocytes treated with rPMD or in combination with PMIs were smeared and air dried on a clean grease free glass slide and imaged using WITec alpha using NSG30 probes with force constant of 22-100 N/m, resonant frequency of 240-440 Hz, tip curvature radius of 10 nm (Tips nano) in non-contact mode. Topographic images were obtained at points per line and lines per image of 512*512 with the scan rate of 0.5 times/line (Trace) (s). All the AFM images were recorded using the Control Four 4.1 software. The images were 3D processed and analysed using software Project Four 4.1 software (WITec, Germany).

### Raman imaging and analysis

The Raman measurements of all the samples was performed using WITec alpha 300RA combined confocal raman microscope. The spectrometer was equipped with solid-state diode lasers operating 532 nm and a suitable CCD detector that was cooled to −60 °C. A Zeiss Fluor (100X) EC Epiplan-NEOFLUAR objective was used. The spectral resolution was equal to 1cm^−1^. The integration time for a single spectrum varied from 2 to 5s. Raman measurements and data analysis were performed using software Project Four 4.1 software (WITec, Germany). All Raman spectra presented were after pre-processing (base line correction, smoothening and background removal) using asymmetric least squares smoothing method.

### Statistical analysis

The data for the half maximal inhibitory concentration (IC_50_) value and the % growth inhibition activity of compounds are expressed as the mean ± standard deviation (SD) of minimum two independent experiments done in duplicates. IC_50_ values were analyzed and calculated using non-linear regression in Graph Pad Prism 8 (CA, USA). Student’s t-test was performed to calculate the *p* values, where p<0.05 represents *; p<0.01 represents ** and p<0.005 represents *** significance.

## Acknowledgements

A.S, R.H, R.A, C.B, are supported by the Shiv Nadar University fellowship. S.G is a recipient of DST Inspire faculty award. S.P is grateful for the support from Shiv Nadar foundation. The authors would like to thank AIRF for access to live cell microscopy, AFM and Raman facilities at JNU. This work has been funded by the National Institute of Health [(NIH, Grant no-U19AI089676-09)], United States and National bioscience award from DBT (S.S, Shailja Singh). The funders had no role in study design, data collection and analysis, decision to publish or preparation of the manuscript.

## Author contributions

A.S, S.G, performed the recombinant protein and inhibitors based *in vitro* and *in vivo* assays. S.S (Subhabrata Sen), C.B synthesised and characterized the inhibitors. R.H helped design and perform *in vitro P.falciparum* assays. S.P, A.R. performed the *in silico* experiments and data analysis. S.G, A.S, A.P.S, I.K performed the liver stage inhibition assay. S.S performed the gametocyte inhibition assay. S.S (Shailja Singh), S.G conceived and designed the experiments. A.S, S.G, A.P.S and S.S (Shailja Singh) analysed the data and wrote the manuscript. All the authors commented on the manuscript.

## Supporting Information

**S1 Fig.**
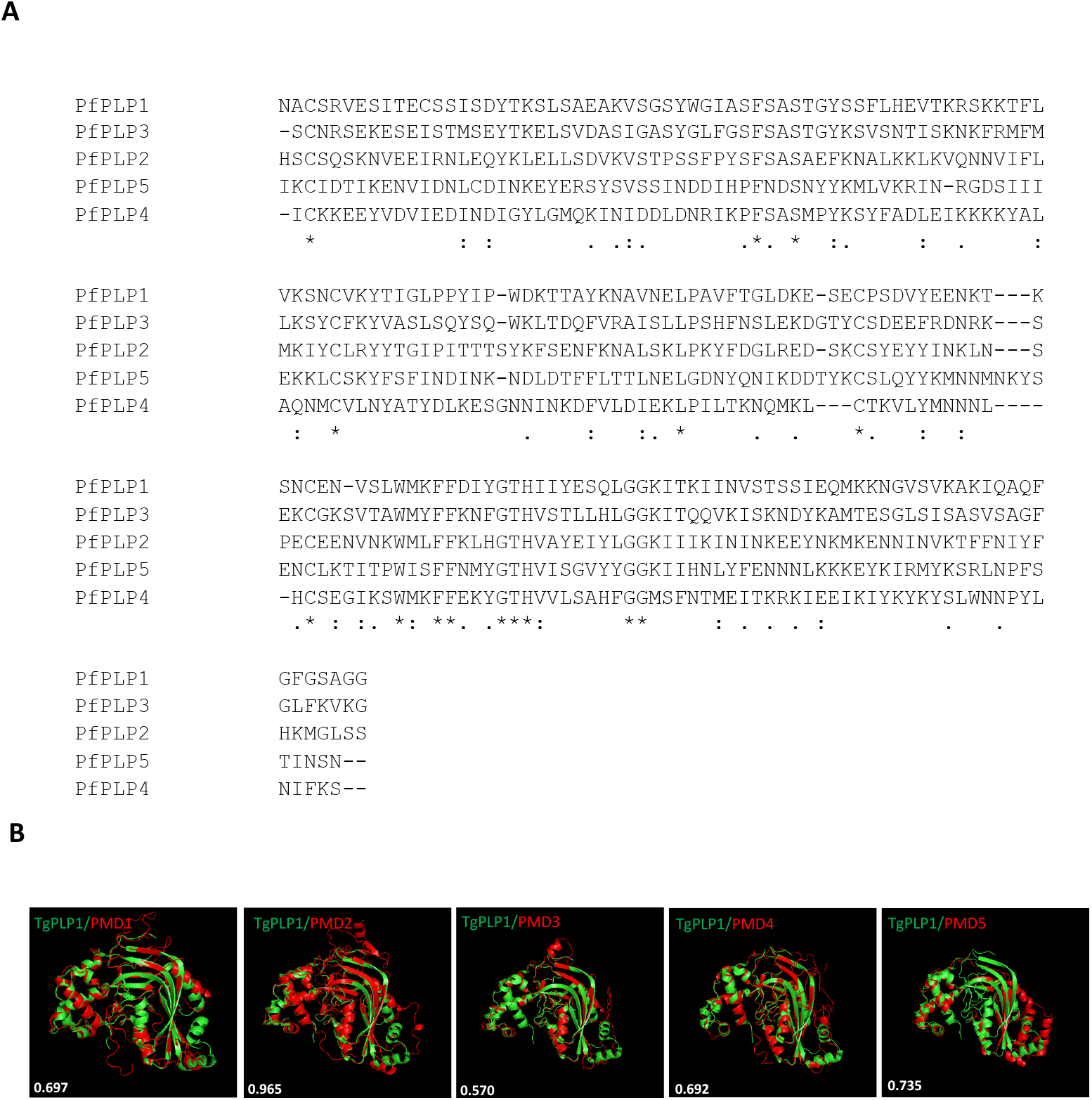
Sequence and structural alignment of the MAC domains of PfPLP1-5. (**A**) The ClustalW alignment of MACPF domains of PfPLP1-5. Conserved residues are highlighted in yellow. (**B**) Structural superimposition of PMDs with MACPF domain of TgPLP1 is shown. The RMSD values are indicated in white.

**S2 Fig.**
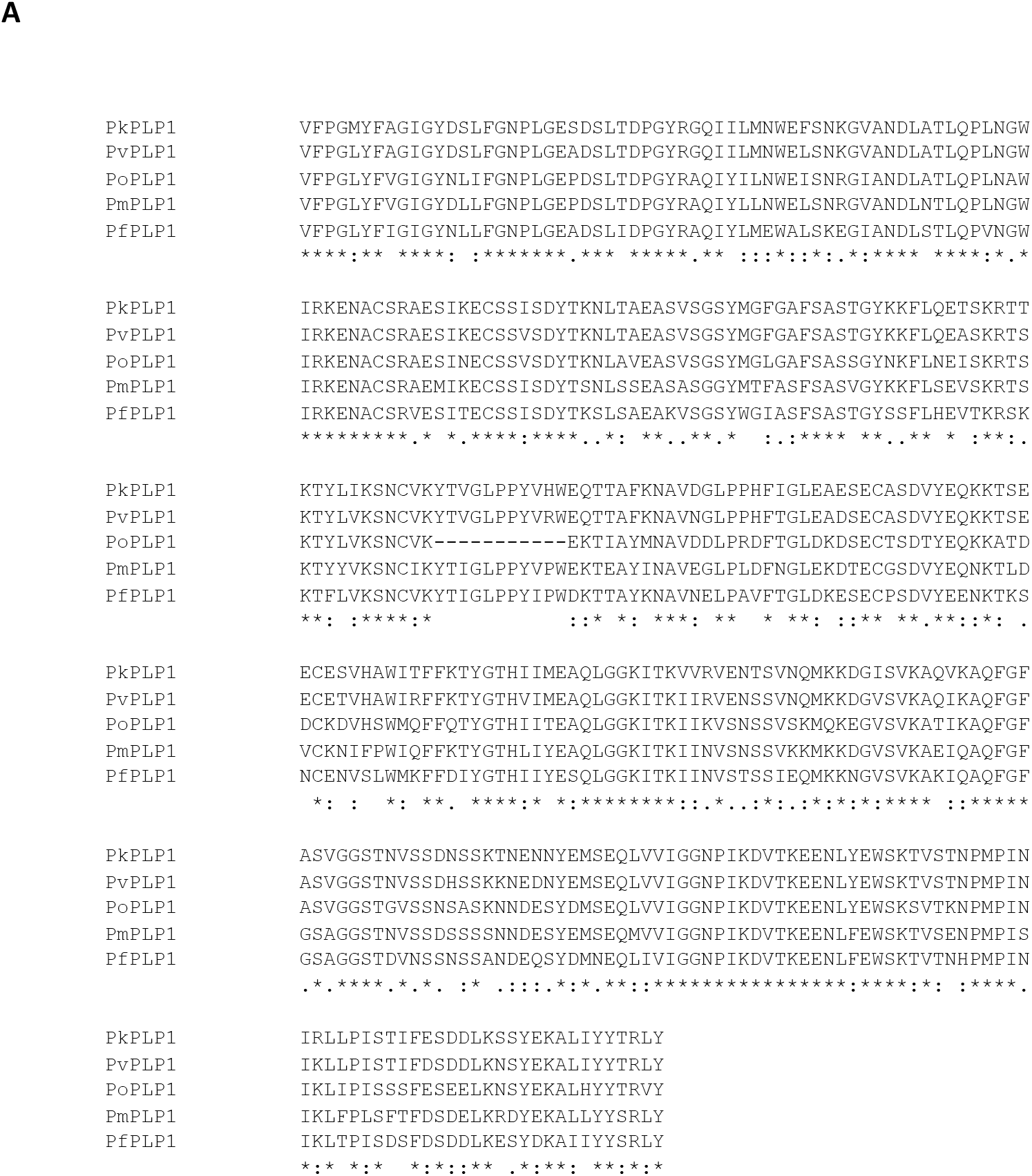
Sequence conservation of MAC domains between different *Plasmodium spp*. (**A**) The ClustalW alignment of MACPF domains of *Plasmodium spp.* PLPs [*Plamodium falciparum* (PfPLP1), *Plamodium vivax* (PvPLP1), *Plamodium ovale* (PoPLP1), *Plamodium knowlesi* (PkPLP1) and *Plamodium malariae* (PmPLP1)] is shown. Conserved residues are highlighted in yellow.

**S3 Fig.**
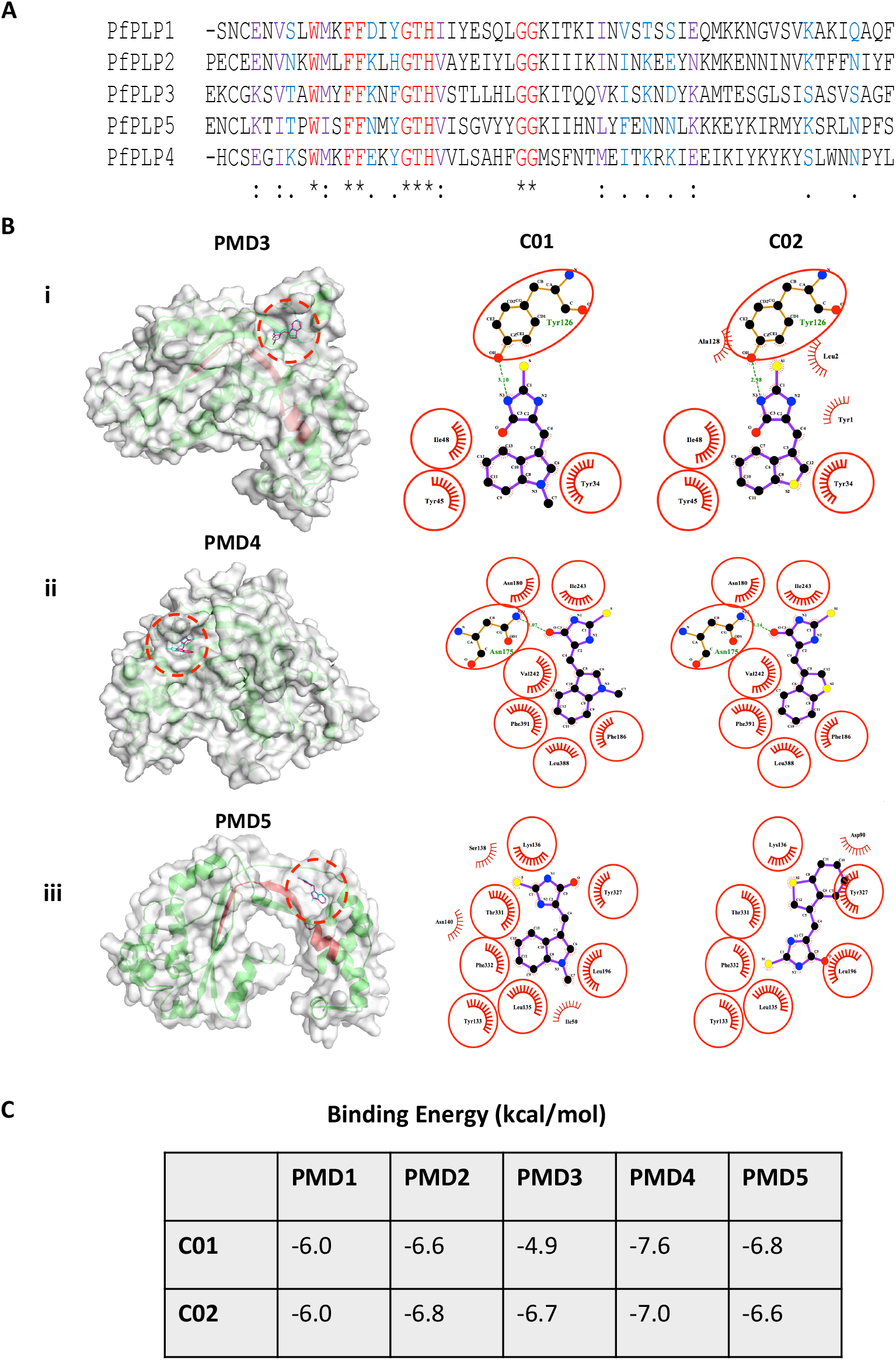
Binding of C01 and C02 to the MACPF domains. (**A**) The ClustalW alignment of the sequences of PMD1-5 where PMIs are binding. Conserved residues are shown in red, blue and purple. (**B**) The surface images of docked complexes showing binding of C01 and C02 to the signature motif of PMD3, PMD4 and PMD5. The ligplot figures demonstrate the specifics of the atoms involved in interaction. (**C**) The table summarizes the binding energies of PMI interaction to PMDs.

**S4 Fig.**
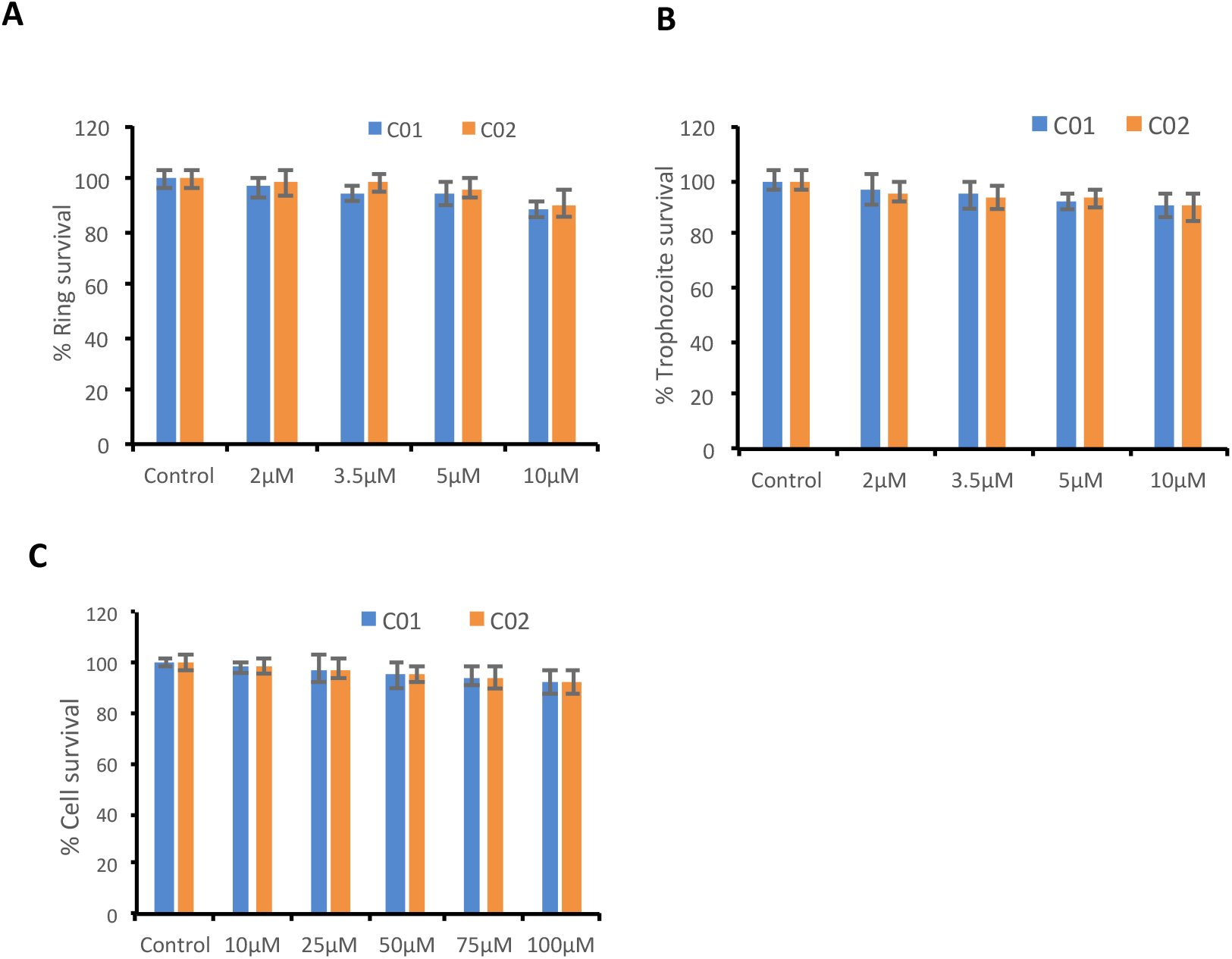
Toxicity analysis of C01 and C02. (**A**) Ring stage parasites were treated with different concentration of compounds for 6 hrs and washed. and the inhibition was accessed by counting the giemsa smears after one cycle of parasite growth. (**B**) Trophozoite stage parasites were treated with different concentration of compounds for 6 hrs and the inhibition was accessed by counting the giemsa smears after one cycle of parasite growth. (**C**) The HepG2 cells were treated with higher concentration of PMIs and toxicity was assessed using MTT assay.

**S5 Fig.**
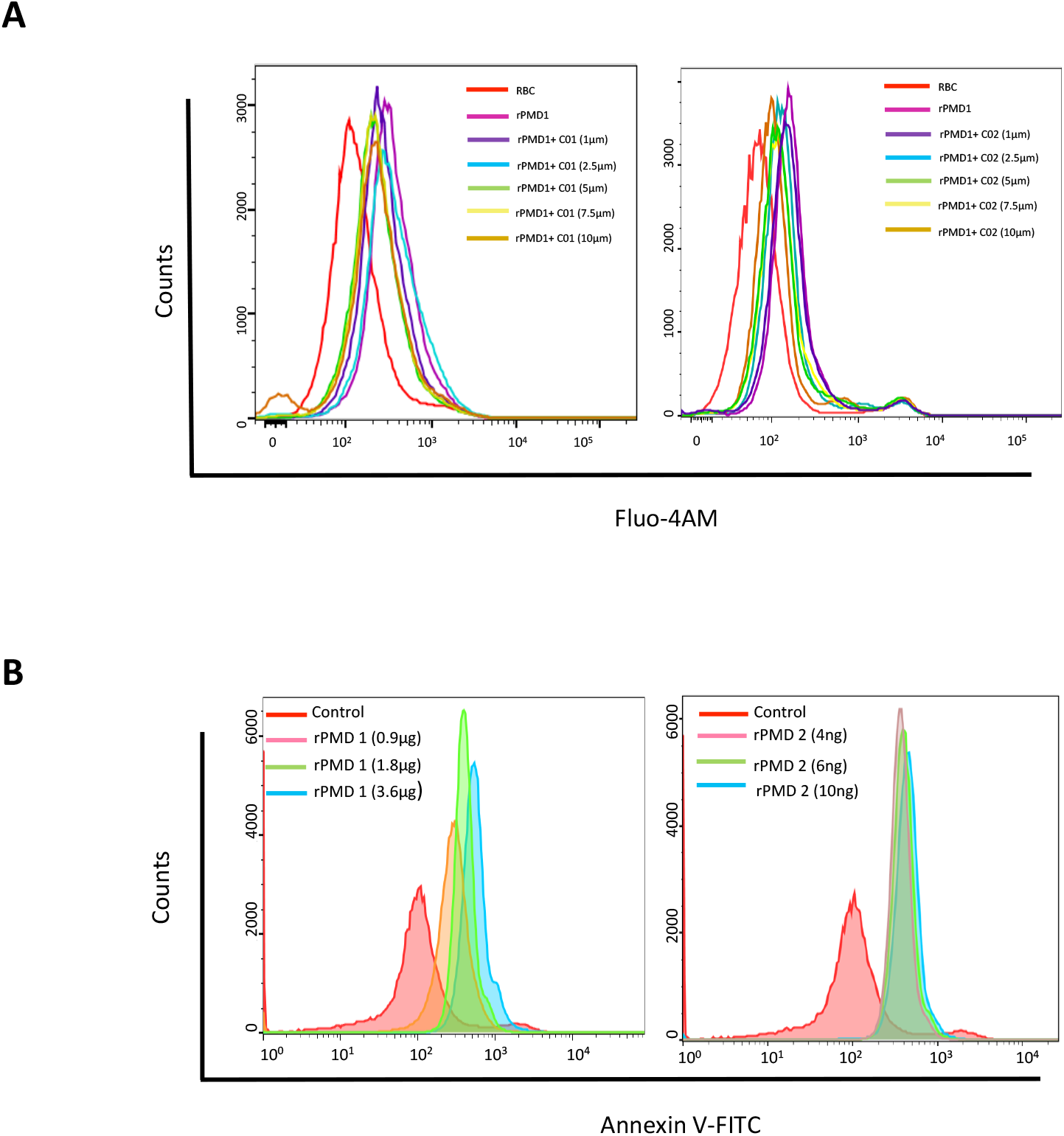
PMI mediated inhibition of Calcium influx and Annexin positivity. (**A**) PMI mediated inhibition in calcium influx was measured by flow cytometry. The Fluo-4AM loaded erythrocytes were treated with rPMD1 in the presence and absence of indicated concentration of PMIs and calcium increase was monitored by Flow cytometer. (**B**) The erythrocytes were treated with rPMD1 or rPMD2 and stained with Annexin V-FITC after 48 hrs. The annexin positive erythrocytes were quantitated using flow cytometer.

**S1 Movie. Live cell imaging of Calcium influx in RBCs upon rPMD2 exposure**

**S2 Movie. Live cell imaging of Calcium influx in RBCs upon rPMD2 and C01 exposure**

**S3 Movie. Live cell imaging of Calcium influx in RBCs upon rPMD2 and C02 exposure**

